# Combination of high pressure treatment at 500 MPa and biopreservation with a *Lactococcus lactis* strain for lowering the bacterial growth during storage of diced cooked ham with reduced nitrite salt

**DOI:** 10.1101/2020.07.22.215863

**Authors:** Stéphane Chaillou, Mihanta Ramaroson, Gwendoline Coeuret, Albert Rossero, Valérie Anthoine, Marie Champomier-Vergès, Nicolas Moriceau, Sandrine Rezé, Jean-Luc Martin, Sandrine Guillou, Monique Zagorec

## Abstract

We investigated the combined effects of biopreservation and high pressure treatment on bacterial communities of diced cooked ham prepared with diminished nitrite salt. First, bacterial communities of four commercial brands of dice cooked ham from local supermarkets, were characterised and stored frozen. Second, sterile diced cooked ham, prepared with reduced level of nitrite was inoculated with two different microbiota collected from the aforementioned commercial samples together with a nisin producing *Lactococcus lactis* protective strain able to recover from a 500 MPa high pressure treatment. Dices were then treated at 500 MPa for 5 minutes and bacterial dynamics was monitored during storage at 8°C. Depending on samples, ham microbiota were dominated by different Proteobacteria (*Pseudomonas, Serratia, Psychrobacter*, or *Vibrio*) or by Firmicutes (*Latilactobacillus* and *Leuconostoc)*. Applied alone, none of the treatments stabilized durably the growth of hams microbiota. Nevertheless, the combination of biopreservation and high pressure treatment was efficient to reduce the growth of Proteobacteria spoilage species. However, this effect was dependent on the nature of the initial microbiota, showing that use of biopreservation and high pressure treatment as an alternative to nitrite reduction for ensuring cooked ham microbial safety merits attention but still requires improvement.

## 1. Introduction

Nitrite salts are used since ancient times as curing agents for the production of cured meat products. Nitrite (and eventually nitrate) salts are commonly added in the brine for the manufacturing of cooked ham. Their role is important for the typical pinky/reddish color development of cured meats [1]. Nitrite salts also participate to the hurdle technology for ensuring microbial safety as bactericidal and bacteriostatic agents against several pathogenic bacteria occurring in meat products, in particular *Clostridium botulinum* [2]. However, nitrite can potentially lead to carcinogenic nitrosamine production in these products, which raises the concern of a potential public health risk [3]. In European Union (EU) all additives authorized before the 20^th^ of January 2009 had to be re-evaluated by 2020. In that context, the European Food Safety Authority (EFSA) re-assessed the safety of nitrite and nitrate salts in 2017. The current acceptable daily intakes for nitrite and nitrate were assessed and experts stated that exposure to nitrites used as food additives was within safe levels for most of the population, except for children [4]. Nevertheless, although experts evaluated the need for further scientific information, a positive link could be established between dietary intake of nitrite (or of processed meat containing both nitrite and nitrate) with some cancers [5, 6]. Therefore, reducing nitrite levels in processed meats appears necessary in order to limit this sanitary issue. Several hurdle technologies can be proposed in order to compensate a higher risk of microbial safety in cooked ham with reduced nitrite salts. Among those, High Pressure Processing (HPP) has been proposed in context of food additive reduction [7 for a review] and was shown to efficiently reduce bacterial contaminants in cooked ham after applying 400-600 MPa treatment for 3-20 minutes depending on the study [8-18] with many reporting an effect at 400 or 500 MPa [9-12, 14-16]. As well, biopreservation, a method using protective cultures, often lactic acid bacteria (LAB) or their metabolites, was described more than 20 years ago for fighting undesired bacterial contaminants [19]. Addition of such LAB protective cultures to cooked meat products, and in particular cooked ham was indeed reported [20, 21]. HPP combined with biopreservation has also been studied [22]. HPP treatment of cooked ham treated with bacteriocins has been shown to reduce the population of several bacterial pathogens or to prolong cold storage [12, 13, 16, 23, 24]. In the present study, we investigated the combined effect of HPP and biopreservation on the dynamics of bacterial communities of cooked ham with reduced level of nitrite salts. We had previously selected a nisin producing *Lactococcus lactis* strain which, although sensitive to HPP treatment, was able to recover and regrow after a 500 MPa HPP treatment [25]. Here, we first determined and collected bacterial communities present on commercial diced cooked ham. Then the dynamics of these bacterial communities was monitored, following their inoculation in sterile diced cooked ham with reduced nitrite level, in the presence of the bioprotective strain, and after the application of the HPP treatment. Our study provides data in link with the following questions: does HPP display a generic effect on microbial inactivation or is there a microbiota signature in pressurized ham? Is the use of nisin-producing *L. lactis* mediated biopreservation a worth hurdle strategy when combined with HPP, and, is this biopreservation effect also microbiota dependent?

## 2. Materials and Methods

### 2.1. Bacterial strains, media and growth conditions

*Lactococcus lactis* CH-HP15 [25] was first cultivated on M17 agar plates (Biokar diagnostic, Beauvais, France) at 30°C for 72 h. One colony was inoculated for preculture in Medium Modelling Ham (MMH) [26] at 30°C for 24 h, and then cultured in MMH at 30°C under agitation at 65 rpm for 13 h until early stationary phase. Bacterial enumerations were performed by plating serial dilutions of bacterial cultures, microbiota or ham stomached samples. The total viable counts were determined on Plate Count Agar (PCA) (Biokar diagnostic, Beauvais, France). Psychrophilic and mesophilic counts were determined after incubation for 4 days at 15°C or 24 h at 30°C, respectively. LAB were enumerated after 4 days incubation at 25 °C on de Man Rogosa and Sharpe (MRS) agar medium pH 5.2 (AES, France) containing bromocresol green (25 mg·l−1) to estimate LAB diversity as previously described [27]. In ham samples inoculated with *L. lactis* CH-HP15, M17 agar plates were used for enumeration after 48 h incubation at 30°C. For anaerobic conditions, plates were incubated in jars with anaerocult sachets (Anaerocult A, Merck, Germany).

### 2.2. Ham sampling and microbiota recovery

Diced cooked ham (except for one sample consisting of sliced cooked ham) was purchased in 2015 and 2016 from local supermarkets in Nantes, France. They were purchased as early as possible considering the use-by-date, transported at cold temperature to the laboratory and reconditioned immediately. These samples were sold as ready-to-eat packs of 120 to 200 g conditioned under modified atmosphere without any indication of the gas mixture used in the packs. At arrival in the laboratory (day 1), bags were opened and dices were immediately reconditioned as small (25 g) aliquots that were further used for bacterial enumeration. Since we had no indication about the gas mixtures of the commercialized samples, we chose to recondition aliquots under air or vacuum packaging to potentially enrich bacterial diversity and incubated them at 4°C and 8°C. Total counts and LAB counts were enumerated at day 1, 7, 14, 21, and 28. When PCA mesophilic counts reached about 7 log_10_ CFU per gram of ham, microbiota were collected by mixing 25 g diced cooked ham in 75 ml peptone salt (AES, France) for 2 min in a stomacher (Masticator, IUL Instruments, England). The homogenate was filtered through the bag filter and centrifuged through a filter from Nucleospin Plant II Midi kit (Macherey Nagel, EURL, France) at 8000 × *g* during 10 min at room temperature. The bacterial pellet was resuspended in 30 ml peptone salt and aliquoted as 1 ml with glycerol 15% and stored at -80°C. In total, microbiota from ten different cooked ham samples were recovered from dices conditioned either under air or vacuum.

### 2.3. Challenge tests

Low-contaminated diced cooked ham containing 18 g/kg sodium chloride and reduced level of nitrite (25 mg/kg; recommended max. level is 120 mg/kg) were used for challenge tests. Those were produced as previously described [25]. Briefly, pork ham muscles were defatted, trimmed and minced using a 20 mm grid. One kilogram of this grinding was mixed under vacuum with 100 g of brine containing water 74.1 g, nitrite salt 4.6 g (0.6%), sodium chloride salt 15.2 g, sodium ascorbate 0.6 g, and dextrose 5.5 g. The mix was melded As follows: components were mixed together in the brine water by stirring with a whisk, until homogenization. Then the brine was added to the meat by mixing and not by injection and then the mix was vacuum-packed. Hams (2.5 kg) were cooked in a 100% humidity atmosphere (90 min at 55 °C, 60 min at 60 °C and 235 min at 67 °C), cooled at ambient temperature (18 °C) for 20 min and then stored at 3 °C. Ham cubes of 1 cm × 1 cm were prepared aseptically, aliquoted in 100 g portions, and stored vacuum-packed at −20 °C until use. Cooked ham dices were defrosted at 4 °C for 24 h, first inoculated with microbiota and then with *L. lactis* fresh culture. Inoculation was performed by adding the microbial suspensions to ham dices placed in bags and hand mixed during two minutes. Two different microbiota were inoculated in low-contaminated diced cooked ham. Each challenge test was performed twice (two independent experiments). Microbiota previously collected were defrosted rapidly at room temperature, diluted to a final concentration of 6 log_10_ CFU.mL^-1^, inoculated at 4 log_10_ CFU g^-1^ in diced cooked ham (1% v/w inoculation rate) and stored overnight at 4°C for allowing bacterial recovery. *L. lactis* CH-HP15 inoculation was then performed with a *L. lactis* fresh culture as follows. When early stationary phase was reached, the culture was centrifuged. The bacterial pellet was rinsed in a sterile solution of NaCl 0.9% and resuspended at 9.3 log_10_ CFU g^-1^ and 0.5 ml were inoculated in 100 g dice cooked ham for an initial concentration of 7 log_10_ CFU g^-1^. Diced cooked ham inoculated with microbiota but without *L. lactis* were also used as controls.

### 2.4. HP treatments

Cooked ham dices were aliquoted as 20 g portions, vacuum-packed and High-Pressure treated at 500 MPa for 5 min. This combination of pressure intensity and duration was chosen according to our previous study [25] for ensuring survival and further regrowth of *L. lactis* CH-HP15 cells essential for their protective effect. Samples were treated using high pressure in a 3 L vertical high pressure pilot unit (ACB, Nantes, France) equipped with a temperature regulator device. The pressure transmitting fluid used was distilled water. The samples were inserted into the vessel (whose internal temperature was regulated to 20 °C) and processed at a compression rate of 3.4 MPa/s until the targeted pressure was reached. The pressure level was held for 5 min, and decompression was nearly instantaneous (less than 2 s). Water temperature reached 24 °C at 500 MPa because of the adiabatic heating. Once treated, packs were then stored at 8 °C and one portion was used at day 1, 5, 12, 30 for bacterial enumeration and bacterial pellet collection for further DNA extraction. Unpressurized samples were also included as controls.

### 2.5. DNA preparation and amplicon sequencing

Total genomic DNA was extracted from the bacterial pellet as described previously [28] using the PowerFood™ Microbial DNA Isolation kit (MoBio Laboratories Inc., Carlsbad, USA) and the High Pure PCR Template Preparation kit (Roche Diagnostics Ltd, Burgess Hill, West Sussex, UK). For each sample, both DNA extracts were pooled. Then, this DNA sample was used as template for three independent amplifications using either the 16S V3-V4 region of the rRNA encoding gene or an internal 280 bp fragment of the Gyrase B subunit encoding gene *gyrB*, as described previously [28]. All PCRs were performed in triplicate. Replicates were pooled and the amplified DNA was purified with a QIAquick kit (Qiagen, Hilden, Germany). Amplicon size, quality, and quantity were checked on a DNA1000 chip (Agilent Technologies, Paris, France). The MiSeq Reagent Kit v2 (2×250 paired-end reads, 15 Gb output) was used according to the manufacturer’s instructions for library preparation and sequencing on a MiSeq 2 (Illumina, San Diego, USA). The quality of the obtained sequences was checked with FastQ files generated at the end of the run (MiSeq Reporter Software, Illumina, USA) and additional PhiX Control. The corresponding pairs of sequences were then attributed to their respective samples using the individual multiplexing barcodes. Quality controls indicated a PHRED quality score of at least 30 for 94% of the reads and a median number of 65,520 ± 14,180 reads per sample and 95,860 ± 11,300 reads per sample for 16S rDNA and *gyrB* amplicons, respectively.

### 2.6. Operational Taxonomic Unit (OTU) analysis and accession numbers

For each sample, paired-end sequences were then loaded in the FROGS (Find Rapidly OTUs with Galaxy Solutions) pipeline [29], checked for quality and assembled. We retained merged sequences with a size of 280 ± 50 bp for *gyrB* and 450 ± 50 bp for 16S rRNA gene. SWARM clustering [30] was applied on the assembled sequences using a maximal aggregation distance of three nucleotides for 16S rRNA gene sequences; for *gyrB* sequences, clustering was more stringent, with a maximal aggregation distance of two nucleotides in order to potentially assign OTUs to the subspecies level. After chimeras and spurious OTUs (low-abundance and low frequency OTUs arising from sequencing artifacts) removal as described previously [28], the dataset comprised 69 OTUs for the 16S rDNA dataset and 252 OTUs for the *gyrB* dataset. Taxonomic assignment of 16S rDNA OTUs was performed with the Blastn+ algorithm [31] on the SILVA 128 SSU database [32], using a threshold of 97% identity for species assignment. For *gyrB* OTUs, it has been previously demonstrated that PCR amplification of *gyrB* can also recover the paralogous *parE* gene, which encodes the β subunit of DNA topoisomerase IV. In order to determine which of the two genes had been amplified for each species, the OTU sequences were blasted against the *gyrB*/*parE* databases established by Poirier and colleagues [28]. Both *gyrB* and *parE* OTUs were retained in our diversity analysis. We assigned taxonomy to the species level when 95% of a sequence matched over 90% of length coverage found in the database. The last step consisted in comparing the taxonomic assignments obtained from the 16S rDNA and the *gyrB* amplicon sequencing. The taxonomic assignment of the 16S rDNA-based OTUs were then homogenized and improved by those obtained from the *gyrB* analysis. This strategy was particularly applied to 16S rRNA gene OTUs with no clear assignment to the species-level (several species included at the threshold of 97% identity).

### 2.7. Beta-diversity and statistical analysis

Bacterial diversity analyses were performed using the phyloseq package (v1.24.2) of R [33]. For each of the samples, analysis of diversity was either performed at the OTU level or at the species level. In studies at the species level, OTUs with similar taxonomic assignment (both *gyrB* and 16S rDNA) were merged using the TAX_GLOM function of phyloseq package, and their abundance (number of reads) was averaged. This resulted into a dataset of 56 species with different taxonomic assignment. Similarly, comparative analysis between ham samples with different experimental conditions was performed after technical (16S rDNA/*gyrB*) and biological (biological repetition) data were merged with the phyloseq function MERGE_SAMPLES. In this case, species abundance was averaged between the four replicates. Bacterial diversity was compared among different groups of samples with permutational ANOVA, specifically using the adonis function within the vegan package [34].

## 3. Results

### 3.1. Diced cooked ham selection for microbiota recovery

Preliminary tests were performed on diced cooked ham of different brands collected from local supermarkets. Those were purchased as close as possible to the production date, based on an average shelf life of 20-25 days. They were then incubated under vacuum or air packaging at 4°C or 8°C. We observed a large variation of total viable counts between samples, ranging from 2 to 7 log10 CFU·g^-1^ one day after purchase *i*.*e*. 2-7 days after production. No strong influence of packaging was observed on psychrophilic and mesophilic total counts, or MRS counts. Enumeration on MRS plates containing bromocresol green enabled us to estimate the putative diversity of LAB species present in dices. Only one to 3 types of colonies were detected indicating a poor diversity among the most dominant LAB species.

Since our objective was to recover microbiota for further re-inoculation on cooked diced ham before HPP treatment, we aimed at collecting standardized quantities of bacteria, as diverse as possible, and enough DNA for further amplicon sequencing. For that purpose, we designed the sampling as follows: i) diced cooked ham were sampled from supermarkets as close as possible to their production date; ii) immediately after arrival in the laboratory dices were reconditioned under vacuum packaging and under air to eventually increase the bacterial diversity to be collected; iii) packs were then stored at 4°C and analysed every 7 days; iv) when total viable counts reached about 7 log_10_ CFU·g^-1^, microbiota were collected and stored for further analyses and DNA was extracted for amplicon sequencing. Through this strategy, diced cooked ham samples from four different brands out of ten (further referred as HAM_A to HAM_D samples) were obtained with enough bacterial diversity and DNA quality (Table 1). No significant difference in population level was observed between air and vacuum packaging, except in HAM_B for which both PCA and MRS counts were about 1 log higher (respectively 7.97 *vs* 6.53 log_10_ CFU·g^-1^ and 7.74 *vs* 6.82 log_10_ CFU·g^-1^) after storage under air.

**Table 1.**
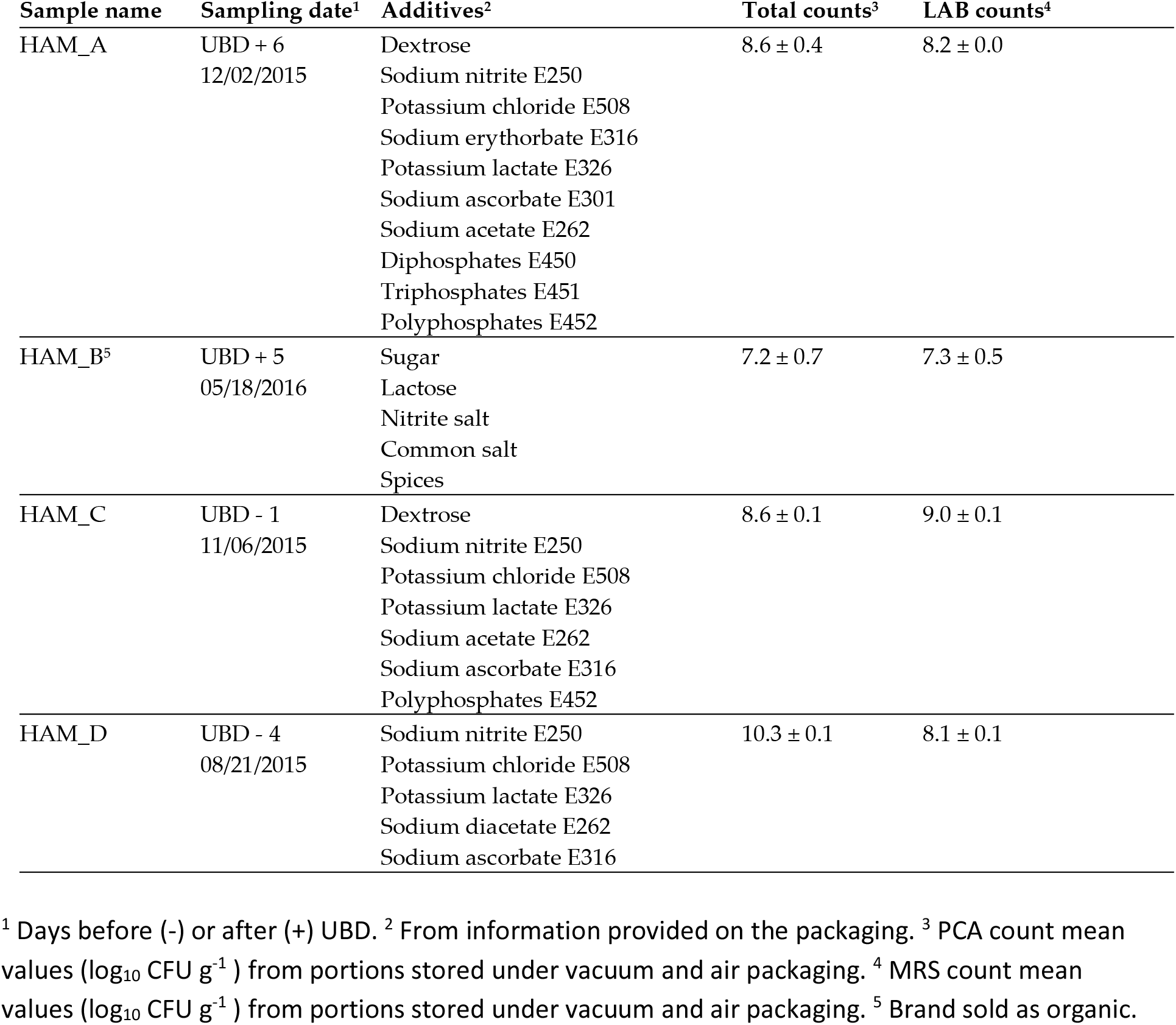
Description of ham specificities.

### 3.2. Four different diced cooked ham can be distinguished by the diversity of their bacterial communities

The bacterial diversity was estimated using a combination of two amplicon sequencing strategies, one using V3-V4 region of the 16S rRNA gene and another one using an internal ∼280 bp fragment of the gyrase B subunit encoding gene (*gyrB*) in order to improve taxonomic assignment of the OTUs to the species or sub-species level as described previously [28]. These two strategies were also used as technical replicates for ensuring that no biases were obtained in the characterization of the four different microbiota as observed in Figure 1A. Only slight differences between both strategies were observed in the clustering of air samples of HAM_C and D. The average number of species detected per sample was 38 ± 5. As shown in Figure 1B, HAM_C and HAM_D samples showed a highly similar microbiota dominated by Firmicutes, in particular *Latilactobacillus sakei* and *Leuconostoc carnosum*, two species commonly found on this type of cured meat products [35-37]. Furthermore, storage type under vacuum or air packaging did not influence significantly the overall abundance of the identified species. On the other hand, the bacterial communities characterized from samples of HAM_A and HAM_B were significantly different from each other and from the aforementioned products, in particular with a clear dominance of Proteobacteria over Firmicutes (Figure 1C). Ham_A microbiota was mainly composed of *Pseudomonadaceae* and *Enterobacteriaceae* with *Pseudomonas lundensis* and *Serratia grimesii* being the most abundant species for Proteobacteria and *Carnobacterium maltaromaticum* being the most abundant species for Firmicutes. Samples from Ham_B revealed to contain the most original microbiota among the four types of ham with the three most abundant 16S rDNA or *gyrB* OTUs being assigned to unknown species as the identity percentage of these OTUs to *gyrB* gene in databases was lower to the ANI (Average Nucleotide Index) threshold of 95% classically used for bacterial species boundary [38]. The most abundant Firmicute *gyrB* OTU was assigned to a *Carnobacterium sp*. showing only 89.0% identity (as best match) to *gyrB* gene of *Carnobacterium funditum* strain DSM 5970. Among Proteobacteria, one OTU was assigned to a *Psychrobacter sp*. with closest match was 89.9% to *gyrB* gene of *Psychrobacter cryohalolentis* FDAARGOS_308, a strain isolated from Siberian permafrost (GenBank: GCF_002208775.2). For the second Proteobacteria OTU, the best identity score found was 91.6 % with *Vibrio sp. SM1977 gyrB* gene, a species isolated from coralline algae surface (GenBank: CP045699.1). It should be noted that the abundance of these two uncharacterized species (*Psychrobacter sp*. and *Vibrio sp*.) varied significantly according to the packaging type used for storage, which may correlate with their 10 times higher counts on air stored samples (Table 1), although we have no evidence for their cultivability in our growth conditions.

**Figure 1.**
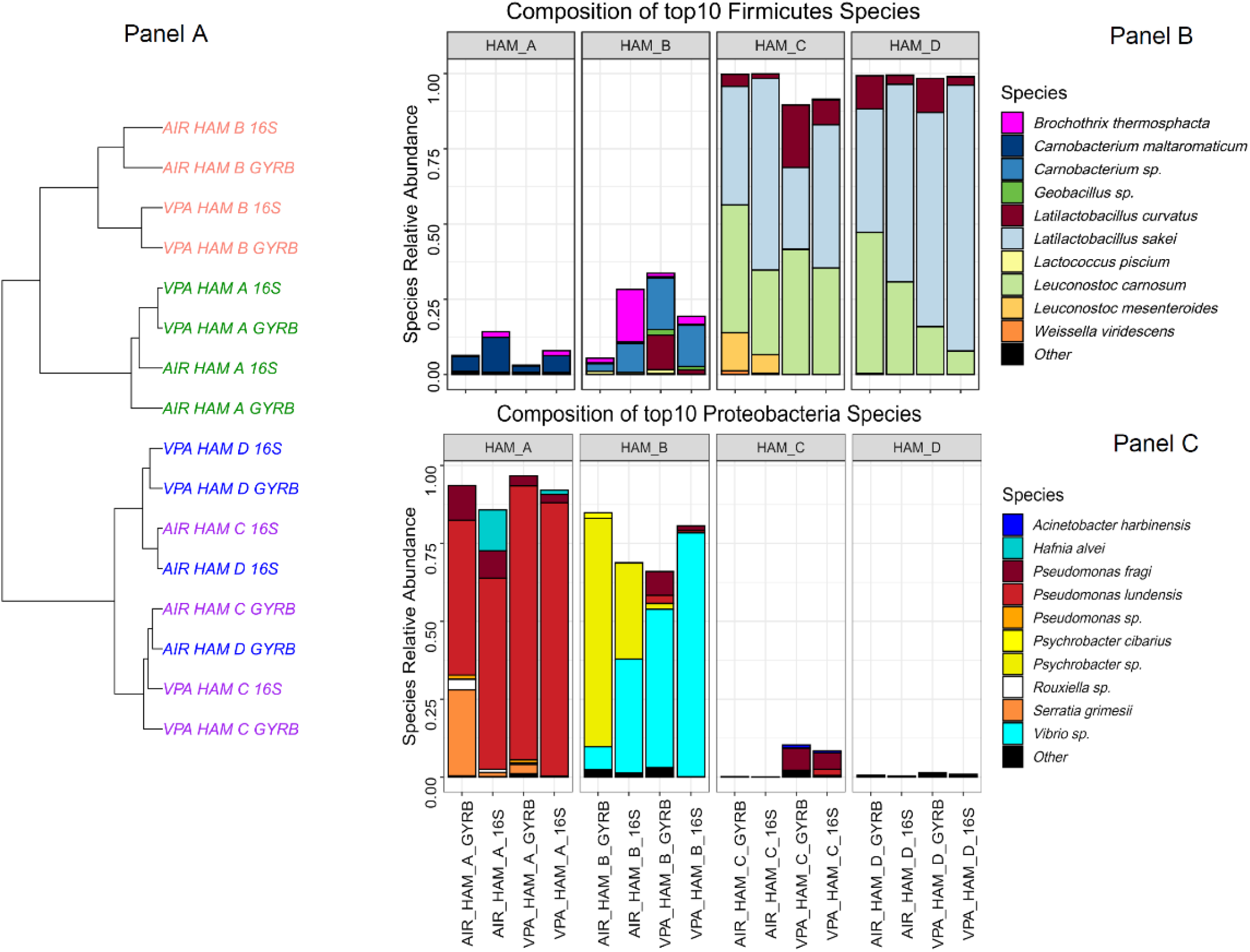
Comparative bacterial community composition between the four brand of cooked ham analysed. Panel A: Cooked ham samples unsupervised clustering tree based on Bray-Curtis distance and Ward algorithm. Samples are colored according to the cooked ham brand. Both air (AIR) and vacuum packed (VPA) samples as well as 16S rDNA-based or *gyrB* –based analysis are shown. Panel B and C show a bar plot composition of the top 10 species identified among Firmicutes and Proteobacteria phyla, respectively. Species are plotted according to their relative abundance in percentage of the whole microbiota. Novel genus nomenclature was applied for *Latilactobacillus* species (*L. sakei* and *L. curvatus*) as recently proposed by Zheng and colleagues [39].

### 3.3. Combined effect of HPP and biopreservation on bacterial community dynamics

For the further steps of our analysis dedicated to monitoring the effect of HPP and biopreservation by *L. lactis* CH-HP15 on the ham microbiota dynamics during storage at 8°C, we decided to focus on the two different microbiota from HAM_A and HAM_B. This was based on the fact that these two microbiota showed a community structure enriched in bacterial species with highly spoilage potential (Proteobacteria) in comparison to microbiotas of HAM_C and HAM_D enriched in *Latilactobacillus*; those latter species being generally rather considered as positive microbial component [20, 21]. In addition, we chose HAM_B microbiota because of the unexpected presence of the two uncharacterized *Psychrobacter* and *Vibrio* species. Table 2 is summarising the estimated bacterial population level at different storage times after the several processing treatments. Similarly, Figure 2 is depicting the relative abundance of each species via 16S rDNA and *gyrB* amplicon sequencing.

**Table 2.** Description of ham specificities.

| Population level in log<sub>10</sub> CFU·g<sup>-1</sup> of nitrite-reduced diced cooked ham |  |  |  |  |
| --- | --- | --- | --- | --- |
| Storage time in days<sup>1</sup> |  |  |  |  |
| Sample name | Day 1 | Day 5 | Day 12 | Day 30 |
| HAM_A samples |  |  |  |  |
| No treatment | 4.63 ± 0.09 | 7.32 ± 0.14 | 7.84 ± 0.10 | 8.15 ± 0.15 |
| HPP | 1.20 ± 0.14 | 3.74 ± 0.01 | 3.00 ± 1.00 | 4.02 ± 0.59 |
| *L. lactis* | 7.21 ± 0.62 | 9.18 ± 0.02 | 8.39 ± 0.12 | 8.87 ± 0.87 |
| HPP + *L. lactis* | 2.54 ± 0.45 | 3.97 ± 0.26 | 5.42 ± 1.27 | 7.60 ± 0.03 |
| HAM_B samples |  |  |  |  |
| No treatment | 5.24 ± 0.18 | 6.10 ± 0.13 | 6.90 ± 0.33 | 6.50 ± 0.20 |
| HPP | 1.39 ± 0.09 | ND<sup>2</sup> | 5.33 ± 0.34 | 6.23 ± 0.42 |
| *L. lactis* | 7.25 ± 0.51 | 8.29 ± 0.01 | 7.82 ± 0.21 | 8.50 ± 0.50 |
| HPP + *L. lactis* | 2.75 ± 0.44 | 3.64 ± 0.07 | 7.11 ± 0.13 | 10.2 ± 0.20 |

<sup>1</sup> Population data were calculated and averaged from the two biological replicates and from two plating culture condition enumerating both total mesophilic bacteria and lactic acid bacteria (see M&M). <sup>2</sup> Not Determined.

**Figure 2.**
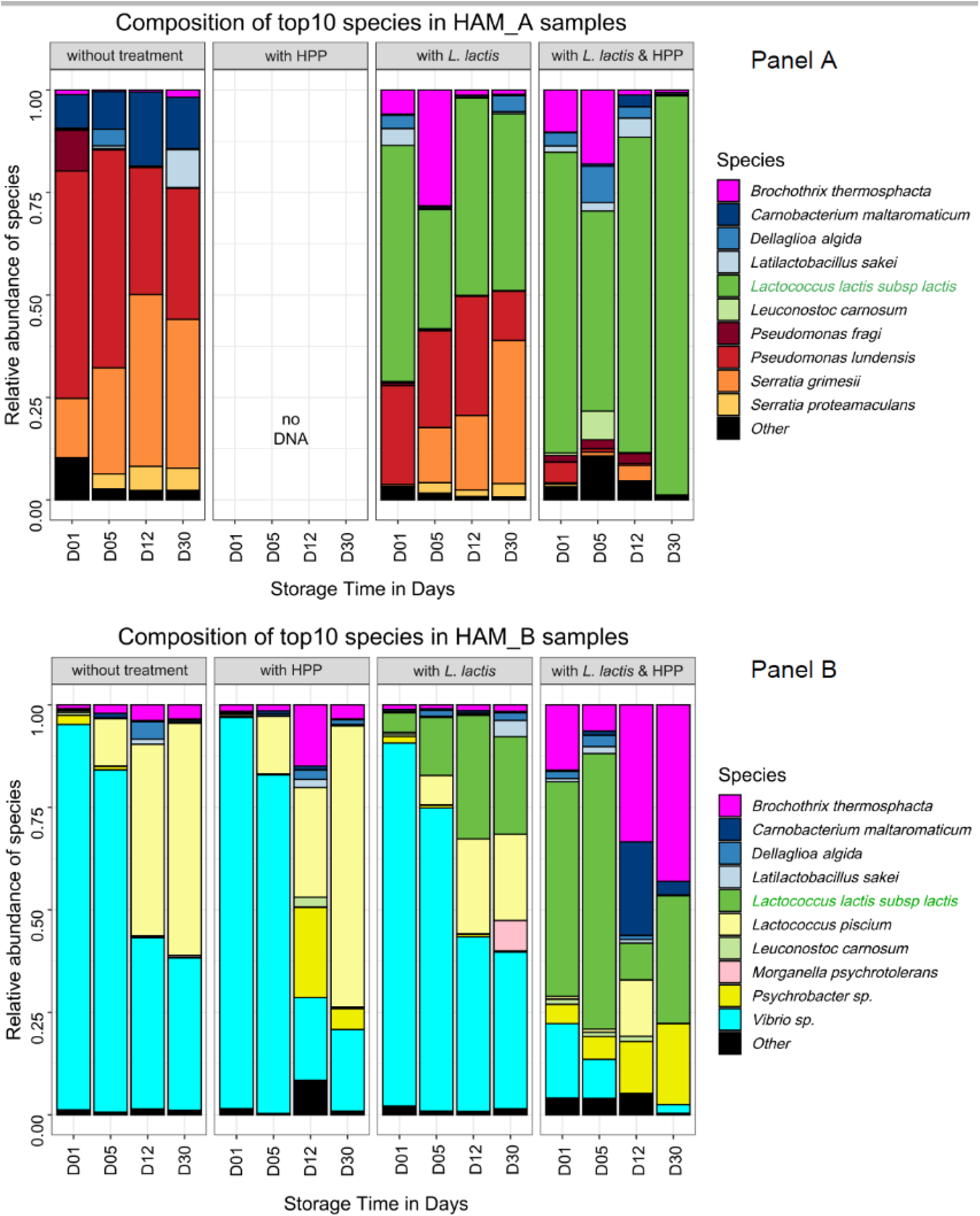
Time course dynamic barplot composition of bacterial communities in nitrite-reduced cooked ham samples upon various treatments. Data for HAM_A and HAM_B samples are shown in panel A and B, respectively. Species are plotted according to their relative abundance in percentage of the whole microbiota. The *L. lactis* strain CH-HP15 used for biopreservation is highlighted in green in the left legend. Each bar plot is the mean of two biological replicates and each replicate value being the average of both 16S rDNA and *gyrB* amplicon sequencing data. As described in Figure 1 for *Latilactobacillus* species, novel genus nomenclature was applied for *Dellaglioa algida* (former *Lactobacillus algidus*) as recently proposed by Zheng and colleagues [39].

The control samples without any processing treatment revealed that at day 1 after inoculation, the microbial communities were similar to the microbiota from HAM_A and HAM_B analyzed in Figure 1, showing that the dices prepared for the challenge-tests were indeed poorly contaminated and that the microbiota had recovered with no strong bias from freezing. HAM_A microbiota grew rapidly under vacuum from ∼4 log_10_ CFU. g^-1^ to reach ∼7 log^10^ CFU·g^-1^ at day 5 and ∼8 log^10^ CFU·g^-1^ at day 30. In contrast, HAM_B microbiota showed a more reduced growth dynamics, reaching at most ∼7 log^10^ CFU·g^-1^ during the whole storage period. Interestingly, the structure of the communities also changed during storage: in HAM_A samples, *P. lundensis*, a species abundant at the beginning of storage was progressively overgrown by *S. grimesii* whereas, *C. maltaromaticum*, the third abundant species, remained at a stable proportion within the community; in HAM_B, the dominant *Vibrio sp*. was also overgrown over time by *Lactococcus piscium*, although this latter species was largely subdominant (<0.1%) in the initial microbiota.

Processing with biopreservation by *L. lactis* CH-HP15 led to two different effects regarding samples from HAM_A and HAM_B. In HAM_A samples, the addition of *L. lactis* had a strong reduction effect on the abundance of Firmicutes species, in particular on *C. maltaromaticum*, which thus became largely subdominant. Interestingly *B. thermosphacta* was temporally more abundant and thus presumably taking benefit from this change (figure 2A). Meanwhile, the inoculation of *L. lactis* had no obvious effect on Proteobacteria species whose pattern was unchanged overtime compared to control. Indeed, after 30 days of storage, while the microbiota had reached a population level above 8.5 log^10^ CFU·g^-1^, *P. lundensis* and *S. grimesii* species displayed a relative abundance of about 50% of the microbiota; the remaining part of the microbiota being covered by *L. lactis*, although this strain had been inoculated thousand-fold higher than the original microbiota. In HAM_B, the pattern of the dominant *Vibrio sp*. was also not affected in comparison to the untreated samples. Indeed, largely dominant at day 1 its relative abundance decreased over time. However, the addition of *L. lactis* induced a possible competition with *L. piscium*, resulting over time into equilibrium of both species. More generally, *L. lactis* CH-HP15 was less competitive in microbiota from HAM_B samples than in microbiota from HAM_A samples.

Whether or not biopreservation was applied, HPP treatment resulted in major changes in the growth dynamics of both microbiota. HPP treatment alone on HAM_A samples reduced drastically the viability of the original microbiota which remained below 4 log^10^ CFU·g^-1^ after 30 days of storage (Table 2). It is likely that bacterial cells were severely injured, indeed no DNA could be recovered from the bacterial pellet despite the use of several extraction protocol trials. Therefore, diversity analysis by amplicon sequencing could not be performed. Nevertheless, samples treated with both HPP and inoculation by *L. lactis* enabled better recovery of DNA and could provide an overview of the HPP treatment on the HAM_A original microbiota.

Proteobacteria species were the most impacted by HPP treatment, whereas *L. Lactis* CH-HP15 could recover rapidly from this treatment reaching progressively a population level above 7.5 log^10^ CFU·g^-1^ after 30 days, and almost a complete domination of the microbiota. Interestingly, as described above on samples only treated with *L. lactis*, the addition of the bioprotective microorganism increased temporally (up to 12 days of storage) the relative proportion of *B. thermosphacta*, thereby indicating that this latter species is quite resistant to HPP treatment. Unlike microbiota from HAM_A samples, that from HAM_B samples recovered rapidly from HPP treatment alone, almost reaching the population level of untreated samples at 30 days (6 log^10^ CFU·g^-1^ in average). The community structure was itself barely affected by the HPP treatment with only a higher proportion of *Psychrobacter* sp. at longer storage time in comparison to untreated samples. Processing with both HPP and *L. lactis* inoculation confirmed these observations as the population level, combining both initial microbiota and *L. lactis*, recovered rapidly (12 days) to 7.0 log^10^ CFU·g^-1^ and raised further to ∼10 log^10^ CFU·g^-1^ at 30 days, which is almost 4 log higher than the level in untreated ham. However, the HPP treatment was more favorable to Firmicutes species in the end, leading to a domination of *L. lactis* CH-HP15 together with *B. thermosphacta*.

## 4. Discussion

The use of an appropriate HPP treatment to preserve perishable food has been the focus of many studies and reviews [22, 40]. HPP is being used to enhance food safety by reducing the microbial development in final products, meanwhile maintaining at best their nutritional and sensorial values to a level acceptable to consumers [17]. However, the effect of HPP on microbial growth dynamic and community structures has not been widely studied and it is not known whether hurdle strategies should be applied in combination with HPP to efficient microbiota inactivation or stabilisation. Indeed, a recent work carried out with a simplified ham microbiota (five species including *Listeria monocytogenes, L. sakei, B. thermosphacta, C. maltaromaticum* and *Leuconostoc gelidum*) showed that HPP treatment is not sufficient to inhibit growth recovery of the microbiota over long storage time [24]. This is pointing out that there is a clear gap in our knowledge on the HPP efficiency towards various microbial communities which may be present on cooked ham. Although the technology could be a very promising strategy to improve the safety of nitrite-reduced cooked ham, it is necessary to investigate whether it should be used as hurdle with the combination of other strategies such as biopreservation [25].

Our strategy was to recover different microbiota to be reusable for performing challenge-tests. As previously demonstrated by Raimondi and colleagues [35], the microbiota of the four types of cooked ham was poorly diverse but highly variable. Some cooked hams with poor diversity were dominated by *L. sakei, Latilactobacillus curvatus* and *L. carnosum* and their microbiota were not considered further as HPP treatments were previously demonstrated to be efficient against the growth of these species [9]. On the other hand, cooked hams from two other producers were characterized by a more diverse microbiota. These microbiota were composed of a quite different mixture of species from Firmicutes and Proteobacteria phyla, including not yet characterized dominant species. Therefore, these two types of microbiota were chosen as model microbiota to test the efficiency of HPP and biopreservation.

Firstly, our results demonstrate that the reduction of viable cells population by HPP is slightly more effective on Proteobacteria species than on Firmicutes species resulting in a small shift of the dominant population from Proteobacteria to Firmicutes when HPP is applied. This finding corroborates many previous observations made by comparing individual species of Gram-negative and Gram-positive pathogenic bacteria resistance to HPP for instance [22 for a review]. Although the cell surface morphology of both types of bacteria can explain such difference, it is interesting to notice that the recovery of cells after HPP is species (and per se microbiota) dependent. Although both microbiota from HAM_A and HAM_B were re-inoculated at the same population level before HPP treatment, that from HAM_A, composed of *P. lundensis* and *S. grimesii* revealed more sensitive (no or almost no recovery) to HPP than that from HAM_B composed of uncharacterised *Psychrobacter* and *Vibrio* species. The species or even strain-dependent resistance to HPP has already been pointed out for Firmicutes, in particular for *L. monocytogenes* and *Staphylococcus aureus* [13]. The underlying mechanisms are not fully understood as HPP resistance and recovery of bacteria also depend on the cell physiology of bacteria present on the food matrix before the treatment. As well, the matrix composition may influence bacterial recovery [41], as was shown for fat content influencing *L. monocytogenes* recovery from thermal inactivation [42]. Furthermore, the capacity of the bacteria to resist HPP treatment should be dissociated from the ability of the bacterial cells to recover and thrive during the storage conditions. Our data are a good illustration of this phenomenon. For instance, *S. grimesii* revealed more competitive than *P. lundensis* to grow during 30 days at low temperature and perhaps under vacuum packaging, leading to a switch of the two species during storage. Similarly, *L. piscium*, a sub-dominant species in original HAM_B microbiota, could outcompete the initial dominant *Vibrio* species.

Another finding from our work is that the *L. lactis* strain used for biopreservation is not competitive towards ham original microbiota. However, we noticed that it has perceivable effect towards the reduction of other Firmicutes species (e.g. *C. maltaromaticum*), perhaps due to the production of nisin. Indeed, this bacterium has been shown in previous studies to harbour sensitivity to this bacteriocin in vacuum packed meat [24, 43]. Albeit the *L. lactis* CH-HP15 strain was inoculated with a level three orders of magnitude higher than the original ham microbiota, the strain was found between 25% to 50% of overall relative abundance in HAM_A and HAM_B, respectively. We previously observed that the inactivation level of this strain after HPP was more important than that of other LAB species, which was compensated by its better ability to recover and rapidly regrow after the treatment [25]. The lack of *L. lactis* competitiveness is probably due to the specific ecology of ham. Indeed, *L. lactis* CH-HP15 was shown able to grow in sterile diced cooked ham at 8°C, reaching 9 log^10^ CFU·g-1 within 5 days with an initial inoculum of 6 log^10^ CFU·g^-1^ [25]. In the present study, the presence of species belonging to the natural microbiota and thus potentially better adapted may explain the lack of fitness of *L. lactis* CH-HP15. Nevertheless, the combination of high level of inoculation of *L. lactis* with HPP treatment leads to an efficient stabilisation of the original cooked HAM_A microbiota, by limiting the growth of *P. lundensis* and *S. grimesii*. Unlike this situation, results were different for HAM_B samples for which the hurdle strategy could not trigger, over the storage time, the outcompetition of *L. lactis* versus the original HAM_B microbiota. Therefore, it can be concluded that for microbiota of HAM_B, the hurdle strategy failed in stabilizing and inactivating the microbial growth.

Furthermore, our results show that *B. thermosphacta* is a species with a very strong recovery dynamics following the HPP treatment. Such observation has already been made by Teixiera and colleagues using the simplified ham microbiota described above [24]. This finding raises the question of HPP treatment benefit on *B. thermosphacta* containing microbiota as this species is a well-known meat spoilage micro-organism [44].

## 5. Conclusions

From our work, we thus conclude that both HPP treatment and *L. lactis*-based biopreservation is strongly microbiota dependent and thus, the value of this strategy requires a specific assessment for each type of cooked ham production. We recommend that HPP treatment should be evaluated not only for pathogenic bacteria but also on putative spoilage bacteria in order to estimate the specific selection of these undesirable micro-organisms, in particular *B. thermosphacta*.

## Author Contributions

Conceptualization, M.Z., S.C., M.C.V. and S.G.; methodology, M.R. A.R. (ham sample and microbiota collection), J.L.M. (nitrite reduced ham preparation), S.G., V.A., and N.M. (challenge-tests), G.C. (DNA extraction, library preparation and amplicon sequencing); validation, S.C. and M.Z.; formal analysis, S.C.; data curation, S.C..; writing—original draft preparation, M.Z. and S.C.; writing—review and editing, M.Z. and S.C.; visualization, S.C.; supervision, M.Z. and S.C., project administration, M.Z.; funding acquisition, M.Z. All authors have read and agreed to the published version of the manuscript.

## Data Availability Statement

The merged (assembled) sequences have been deposited in the Sequence Read Archive database under the accession numbers SAMN13761947 to SAMN13762161, corresponding to BioProject PRJNA599607.

## Acknowledgments

The authors express their gratitude to the INRAE MIGALE bioinformatics facility (MIGALE, INRAE, 2020. Migale bioinformatics Facility, doi: 10.15454/1.5572390655343293E12) for providing computational resources and data storage. We also thank the INRA @BRIDGe platform for carrying out the MiSeq sequencing runs.

